# Monitoring the production of high diffraction-quality crystals of two enzymes in real time using *in situ* dynamic light scattering

**DOI:** 10.1101/2020.01.05.888370

**Authors:** Raphaël de Wijn, Kévin Rollet, Sylvain Engilberge, Alastair G. McEwen, Oliver Hennig, Heike Betat, Mario Mörl, François Riobé, Olivier Maury, Eric Girard, Philippe Bénas, Bernard Lorber, Claude Sauter

**Author notes:** These authors contributed equally to the work (Raphaël de Wijn, Kévin Rollet). Corresponding Author: Claude Sauter.

## Abstract

The reproducible preparation of well diffracting crystals is a prerequisite for every structural study based on crystallography. An instrument called the XtalController has recently been designed that allows the monitoring of crystallization assays using dynamic light scattering and microscopy, and integrates piezo pumps to alter the composition of the mother liquor during the experiment. We have applied this technology to study the crystallization of two enzymes, the CCA-adding enzyme of the psychrophilic bacterium *Planococcus halocryophilus* and the hen egg white lysozyme in the presence of a synthetic chemical nucleant. We were able to i) detect early nucleation events and ii) drive the crystallization system (through cycles of dissolution/crystallization) towards growth conditions yielding crystals with excellent diffraction properties. This technology opens a way to the rational production of samples for crystallography, ranging from nanocrystals for electron diffraction, microcrystals for serial or conventional X-ray diffraction, to larger crystals for neutron diffraction.

## 1. Introduction

Since its birth in the sixties, biocrystallography has been a primary source of structural information, contributing more than 90% of 3D structures accessible in the Protein Databank [1] and remains a central player in structural biology, alongside NMR and CryoEM. Over the last decade, new experimental setups have been introduced that widen its applicability and transform the daily practice of crystal growers and crystallographers. The recent advent of X-ray Free electron lasers (X-FEL) enables the serial femtosecond diffraction (SFX) analysis of micro- and nano-crystals, and offers unprecedented possibilities for time-resolved experiments [2–4]. At the same time, electron microscopes have been highjacked to perform micro-electron diffraction (μED), opening the way to the characterization of nanocrystals using laboratory-based instruments [5–8]. Though, like conventional ones relying on synchrotron or neutron sources, these new crystallographic approaches require crystalline material and call for the development of means facilitating the production of calibrated samples (i.e. nano-, micro-, or macrocrystals) with a size adapted to the radiation (electrons, X-rays or neutrons) and the experimental setup.

Growing crystals of a new biomolecule (protein, DNA, RNA and their complexes) is often a time-consuming task that involves a trial-and-error screening step to find solvent conditions generating promising crystalline or microcrystalline phases. It is followed by an optimization step to improve the quality of one or several crystalline forms and make them suitable for diffraction analysis [9]. However, before the first diffraction test, the evaluation of this two-step process mainly relies on optical microscopy observations. As a consequence, early crystal growth events, including nucleation, nano-crystal or nano-cluster formation that directly impact the final crystallization outcome, remain hidden to the crystal grower. For this reason there is a clear need for a system enabling the preparation and the optimization of crystals under well-defined and controlled conditions, and ensuring reproducible crystalline properties and quality. The concept of such a system emerged in the nineties in the frame of a European research consortium on crystal growth (European Initiative for Biocrystallogenesis) and was developed in the context of the OptiCryst European consortium [10]. The current implementation called XtalController (or XC900; Xtal Concepts GmbH, Hamburg) is composed of a crystallization chamber for a single experiment with precise temperature and humidity control [11]. The composition of an initial drop (volume 5-10 μl) of a solution of the target biomolecule can be modified by the injection of various solutions (such as water, buffer or crystallant) using two piezo injectors spraying 70 pl droplets (**Figure 1**). The sample drop can also be concentrated by evaporation and its composition (i.e. the concentrations of components) is continuously calculated from its weight recorded to ± 1 μg by an ultra-sensitive balance. The instrument also provides diagnostic means to track the drop content along the experiment and to navigate in the phase diagram, from an undersaturated solution to a supersaturated state leading to crystal growth or precipitation. It can detect in real time the early occurrence of association events leading to nucleation, as well as of nanocrystals by dynamic light diffusion (DLS) and the growth of crystals by video microscopy as soon as they reach a size exceeding a few microns. Experimental conditions can be varied and monitored in real time to stabilize a specific phase or drive the system in the phase diagram towards another phase.

**Figure 1:**
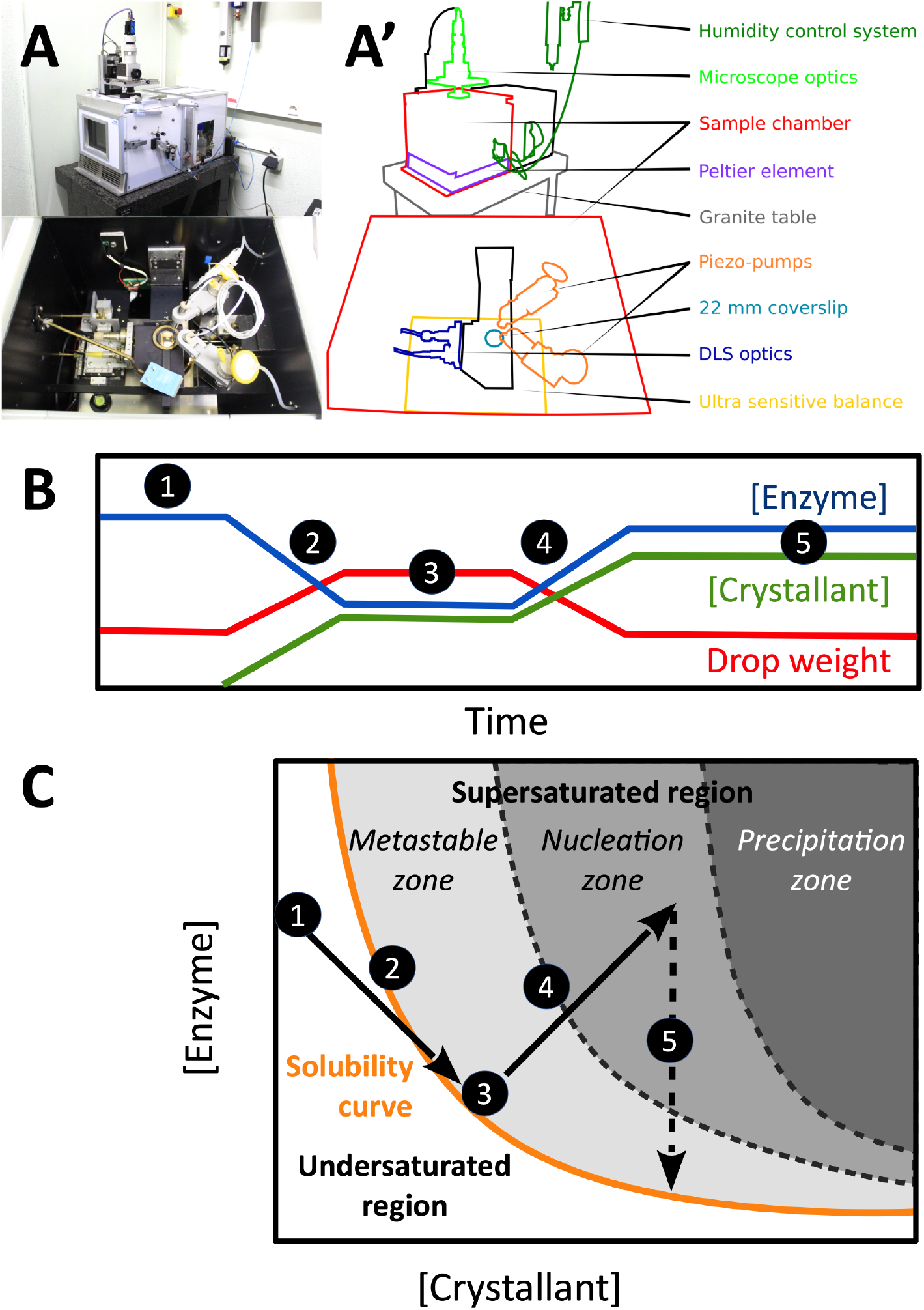
XtalController setup. A) Side and top pictures of the instrument and A’) schematic view highlighting its major components. B) Example of experimental schedule with curves indicating the variation of drop weight (red curve) and associated variations of enzyme and crystallant concentrations (blue and green curves, respectively). The same color code will be used in all figures. C) Corresponding trajectory in a theoretical phase diagram. Step 1: incubation of the enzyme solution at constant concentration. Step 2: addition of crystallant (increase of crystallant concentration, decrease of biomolecule concentration in the drop). Step 3: incubation, constant drop weight (compensation of evaporation by water injection) to keep biomolecule and crystallant concentrations constant. Step 4: controlled evaporation of the drop leading to increased concentrations of biomolecule and crystallant to reach the nucleation zone. Step 5: incubation until crystals start to grow, consume part of the soluble enzyme stock and bring the system back to equilibrium on the solubility curve.

For more than three decades DLS has proven to be instrumental to study nucleation, to perform quality control of biological samples, to predict the propensity of the latter to crystallize, and, more recently, to follow their behavior in crystallization assays [12–16]. With the unique and innovative combination of piezo injectors to modify the experimental conditions and DLS to track in real time the effect of various physical-chemical parameters (chemical composition, biomolecule concentration, temperature) on biomolecules in solution, the XtalController opens a wealth of possibilities for basic and applied crystallogenesis. First examples included the observation of liquid dense clusters formed during nucleation [17] and the preparation of crystals with well defined size [18].

Here we used this technology to study the crystallization of two enzymes, the CCA-adding enzyme from the psychrophilic bacterium *Planococcus halocryophilus* (PhaCCA) and the egg-white lysozyme from hen (HEWL). In the first case, classical vapor-diffusion assays produced numerous small crystals or precipitates. The XtalController helped better define the appropriate crystallant concentration to nucleate and grow large crystals of PhaCCA. In the second case, we used the XtalController to highlight the nucleating effect of a lanthanide complex, the crystallophore Tb-Xo4 [19,20] on HEWL in the absence of crystallant. Both examples illustrate the potential of this technology in crystallogenesis and for the design of protocols to produce calibrated crystalline samples for a variety of crystallographic applications.

## 2. Material and methods

### 2.1 Chemicals and enzyme samples

Chemicals used for the preparation of buffers and crystallization solutions were of highest purity grade. Solutions were filtered on 0.22 μm porosity membranes. PhaCCA (monomer of 420 amino acids, 48 kDa) was produced in *Escherichia coli* cells, purified, concentrated to 5 mg/ml and stored at 4°C in 50 mM HEPES-NaOH pH 7.5, 100 mM NaCl as described previously [21]. HEWL (monomer of 129 amino acids, 14 kDa) was purchased from Seikagaku Japan (Cat. N° 100940), Roche (Cat. N° 10153516103) and Sigma (Fluka Cat. N° 62970-5G-F). It was used without any further treatment and dissolved in water (Roche) or in 10 mM sodium acetate pH 4.5, 40 mM NaCl (Seikagaku and Sigma) at concentrations ranging from 25 to 71 mg/ml. Stock solutions were filtered on a 0.22 μm Ultrafree-MC membrane (Millipore) prior to concentration measurement. The crystallophore Tb-Xo4 used as nucleant for HEWL was synthesized and purified as described [19]. It was dissolved in water to prepare a 100 mM stock solution.

### 2.2 Crystallization in the XtalController

The humidity and temperature of the crystallization chamber of the instrument were set to 99.5% and 20.0°C, respectively, one hour before starting an experiment to ensure the stability of experimental conditions. A drop of 10 μl of enzyme stock solution was deposited on a siliconized glass coverslip (Ø 22 mm) placed on the balance. One pump was loaded with the appropriate crystallant solution and the second with pure water. The crystallization chamber was closed and the protocol started with a 5 min step to monitor drop evaporation and compensate the loss of weight by injection of water to keep the drop weight constant. Several steps of crystallant addition, drop evaporation or dilution were scheduled to explore the phase diagram. The shooting frequency of the pumps was adjusted to vary the slope of concentration variations. DLS measurements and drop image capture were scheduled at regular time intervals (e.g. every 5 to 30 min) to follow nucleation, aggregation or crystal growth events. At the end of the experiment, the coverslip was transferred onto a 24-well Linbro plate for storage and/or incubation.

### 2.3 Standard DLS measurements

In parallel to XtalController experiments, the effect of Tb-Xo4 on lysozyme was recorded using a Nanostar light scattering instrument (Wyatt Technology, Inc.). 10 μL of Sigma lysozyme (71 mg/mL) were transferred into a quartz cell for DLS measurements at 20°C. The drop was covered with 10 μL paraffin oil and the cuvette was sealed with Parafilm™ foil to prevent evaporation. Subsequently 1 μL of a 100 mM Tb-Xo4 stock solution in 10 mM sodium acetate pH 4.5 was added. In control experiments the Tb-Xo4 solution was replaced by buffer. Data were corrected for solvent viscosity and refractive index.

### 2.4 Crystal analysis

Crystals of PhaCCA were analyzed in cryogenic conditions at FIP/BM30A beamline (ESRF, Grenoble, France) using an ADSC Quantum 315r detector. A crystal was soaked for a few seconds in the mother liquor supplemented with 20% (w/v) glycerol, mounted in a cryoloop and flash-frozen in liquid nitrogen. 240 images were collected with a rotation of 0.5° and an exposure time of 1 s per frame. Crystals of HEWL were analyzed at ambient temperature using a Rigaku FR-X diffractometer at the FRISBI platform (IGBMC, Illkirch, France) with an EIGER R 4M detector (DECTRIS) with a 2θ offset of 10°. Several crystals grown with Roche, Sigma and Seikagaku lysozymes were tested and diffracted to up to 1.5 Å resolution but showed rapid decay due to radiation damage at ambient temperature. The exposure time and rotation speed were adapted accordingly to collect a full dataset from a crystal of Seikagaku lysozyme plunged in viscous Parabar 10312 (Hampton Research), mounted in a cryoloop and protected from dehydration using the MicroRT room temperature kit from MiTeGen. 720 images were collected with a rotation of 0.25° and an exposure time of 2 s per frame. Data were processed with the XDS package [22].

### 2.5 Structure determination

The structures of PhaCCA and HEWL were refined in PHENIX [23] using PDB entries 6QY6 and 6F2I (cleared of solvent molecules and ligands), respectively, as starting models for initial rigid body adjustment. Several rounds of refinement and manual inspection in COOT [24] were performed until convergence of R_free_ (calculated using 5% of reflections). The model of PhaCCA was refined using two TLS groups and includes 365 residues (the N-terminal expression tag and the flexible loop encompassing residues 83-94 are not visible), two phosphate ions, three acetate ions and three glycerol molecules present in the mother liquor and the cryoprotection solution. The model of HEWL was refined using anisotropic atomic displacement parameters (ADPs) and includes the full enzyme sequence, one sodium, two chloride ions, a full Tb-Xo4 complex and two additional Tb2+ sites clearly identified in the anomalous density map but for which the ligand was not observed due to low occupancy.

## 3. Results and discussion

### 3.1 Triggering the growth of large crystals of PhaCCA

We recently reported the crystallization of a new tRNA nucleotidyltransferase or CCA-adding enzyme from the cold-adapted bacterium *P. halocryophilus* living in the permafrost [21]. During the process of optimization we observed that the protein had a tendency to produce either clear drops or drops full of small crystals or light precipitate, indicating a narrow nucleation zone. To increase the reproducibility of crystallization, microseeding was systematically used, both in vapor-diffusion or in counter-diffusion assays [21,25]. In the present work the goal was to exploit the XtalController technology to determine appropriate crystallant concentration sufficient for the nucleation of a minimal number of crystals.

**Figure 2** shows a typical crystallization assay with PhaCCA. The experiment starts with a drop of enzyme at 5 mg/ml in its storage buffer. The crystallant (stock solution: 1 M diammonium phosphate, 0.1 M ammonium acetate pH 4.5) is gradually added into the enzyme solution using a piezo pump. The DLS signal clearly indicates the start of protein association around 0.2 M diammonium phosphate with the appearance of objects larger than the initial PhaCCA monomer and increasing in size. This phenomenon is amplified during the step of drop evaporation when both the concentration of crystallant and enzyme simultaneously increase, hence increasing the supersaturation of the enzyme solution. The monomeric form of PhaCCA remains present throughout the experiment, in equilibrium with nano/microcrystalline objects.

**Figure 2:**
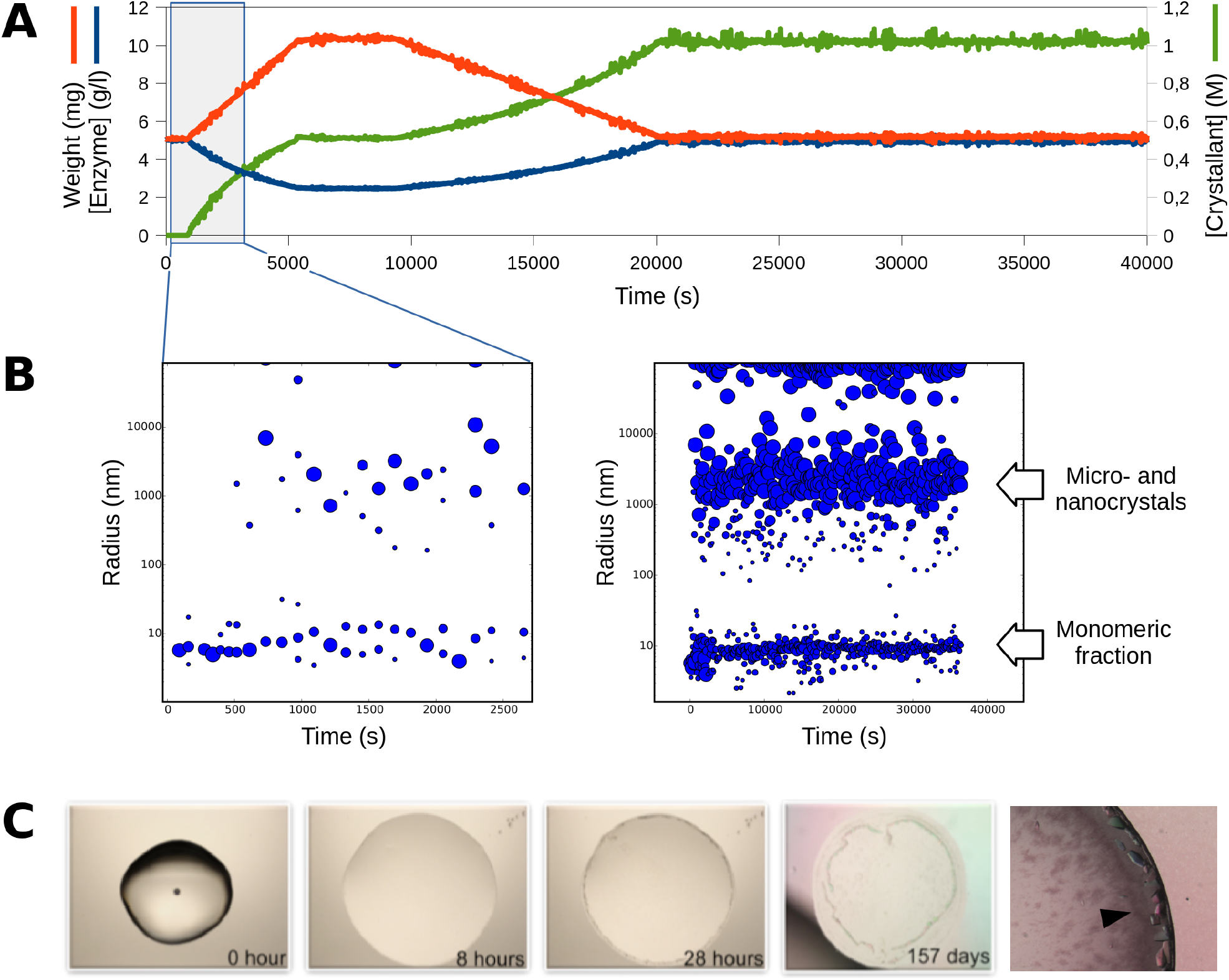
Crystallization of PhaCCA enzyme in the XtalController. A) Experimental parameters are recorded and curves display the variation of drop weight and corresponding variation of enzyme and crystallant concentrations over time. B) The evolution of particle size in solution is monitored by DLS over time. At each time point, different populations detected in solution are represented as blue dots. The center of the dot indicates the hydrodynamic radius *Rh* of the population and the surface its relative contribution to the overall scattered signal. The DLS distribution on the left highlights early events (blue window in A) and shows that the enzyme starts to react at concentrations of crystallant as low as 0.2 M. Particle size distribution on the right corresponds to the monitoring of the complete experiment. The signal of monomeric PhaCCA (*Rh* = 4 nm) decreases while larger objects, likely nano-crystals or nano-clusters, appear upon crystallant addition. C) A selection of drop micrographs taken along the experiment shows the appearance of small objects after 28 hours, leading to useful samples for diffraction analysis within several weeks of incubation over a reservoir in a Linbro plate. The close-up view on the left hand side shows typical PhaCCA crystals used for data collection (**Table 1)**.

After 48h, the monitoring is stopped and the cover slip is transferred from the instrument to a classical 24-well Linbro plate to be stored in equilibrium with a reservoir containing a crystallant solution at the same concentration as that reached at the completion of the protocol (i.e. 1 M diammonium phosphate, 0.1 M ammonium acetate pH 4.5). Small crystals that were already visible after 28 hours in the periphery of the drop, grew slowly to a useful size (100-200 μm) for X-ray diffraction (see 3.3) over a period of several weeks.

In an attempt to increase the size of PhaCCA crystals, protocols including several cycles of drop concentration / dilution were tested. The experimental idea was to trigger protein interaction, then redissolve partially the population of nuclei to select a few for growth in the next drop concentration step. **Figure 3** illustrates this strategy and displays resulting crystals. The largest reached a size of 0.5 mm after 11 days of incubation. This strategy may be exploited in the future to grow large crystals for neutron diffraction. A similar concept was applied using temperature variation coupled with dialysis to adjust the supersaturation and reduce the number of nuclei to promote the growth of large mm-sized crystals [26]. The great advantage of the XtalController is the possibility to monitor the fluctuation of particle populations in real time. The DLS signal shows that the monomer of PhaCCA is converted into larger objects during concentration steps while the drop remains optically clear. If the process is too fast and evolves towards precipitation, the whole system can be driven back to lower concentrations favoring the dissolution of precipitates and the solubilization of monomers. A series of cycles of decreasing concentration variation allows a slow and controlled convergence to the target condition, similar to that used in classical vapor diffusion assays but resulting in many fewer nucleation events and, thus, in larger crystals.

**Figure 3:**
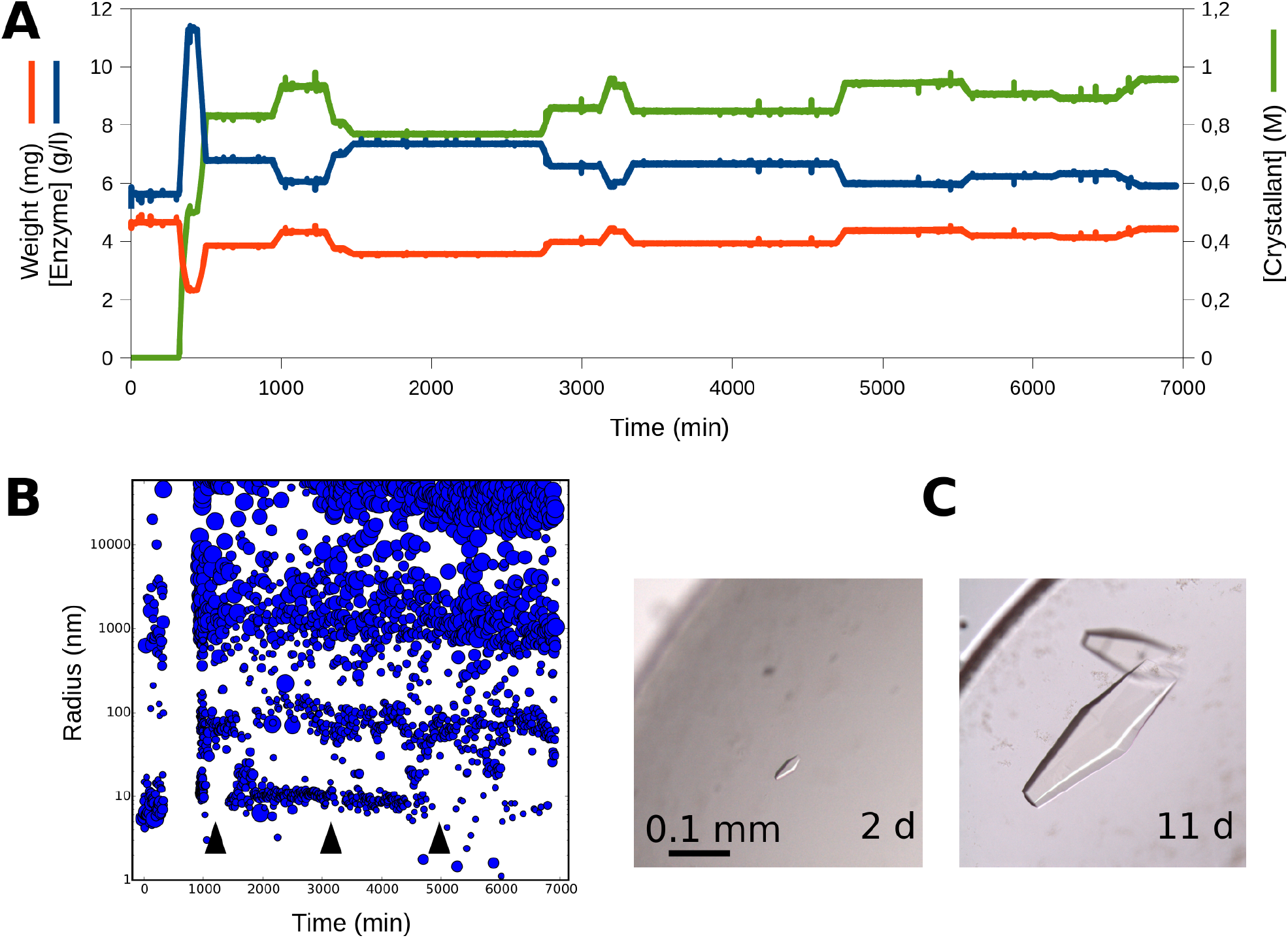
Production of large PhaCCA crystals. A) The crystallant was injected in the sample drop following the same protocol as in **Figure 2.** Four cycles of drop concentration / dilution were performed. B) The DLS monitoring shows that each time the drop is concentrated, the signal for the population of monomeric PhaCCA decreases (see arrows) in favor of larger particles. At the end of this protocol, the clear drop was transferred from the XtalController to a Linbro plate for incubation. First crystals appeared after 2 days and the biggest reached a size 0.5 mm in 11 days.

### 3.2 Tracking the nucleant effect of crystallophore on HEWL

A terbium complex, called crystallophore or Tb-Xo4, with phasing and nucleating properties was recently described [19,20,27]. When preparing lysozyme solutions in pure milliQ water including 10 mM of crystallophore for screening experiments, S. Engilberge observed that the addition of the compound led to the formation of a pellet composed of single lysozyme crystals at the bottom of eppendorf tubes after 20 to 30 days in the cold room. To confirm the ability of the nucleant to promote lysozyme self-association in conditions containing very low salt concentration, the XtalController was used to monitor the behavior of the enzyme in solution upon injection of 10 mM Tb-Xo4 dissolved in water. The control experiment consisted in injecting an equivalent volume of pure water. After incubating 10 μl of a fresh HEWL solution (25 mg/ml) at 20°C to check its stability by DLS (**Figure 4**), the crystallization chamber was opened for a few seconds to inject 1 μl of a 100 mM Tb-Xo4 solution (or water) directly into the drop using a 1 μl Hamilton syringe. Accordingly, the drop weight increased to 11 mg and rapidly went back to 10 mg upon water evaporation controlled by the instrument scheduler. The drop was incubated for hours and DLS measurements were recorded before its transfer to a Linbro plate for long term storage.

**Figure 4:**
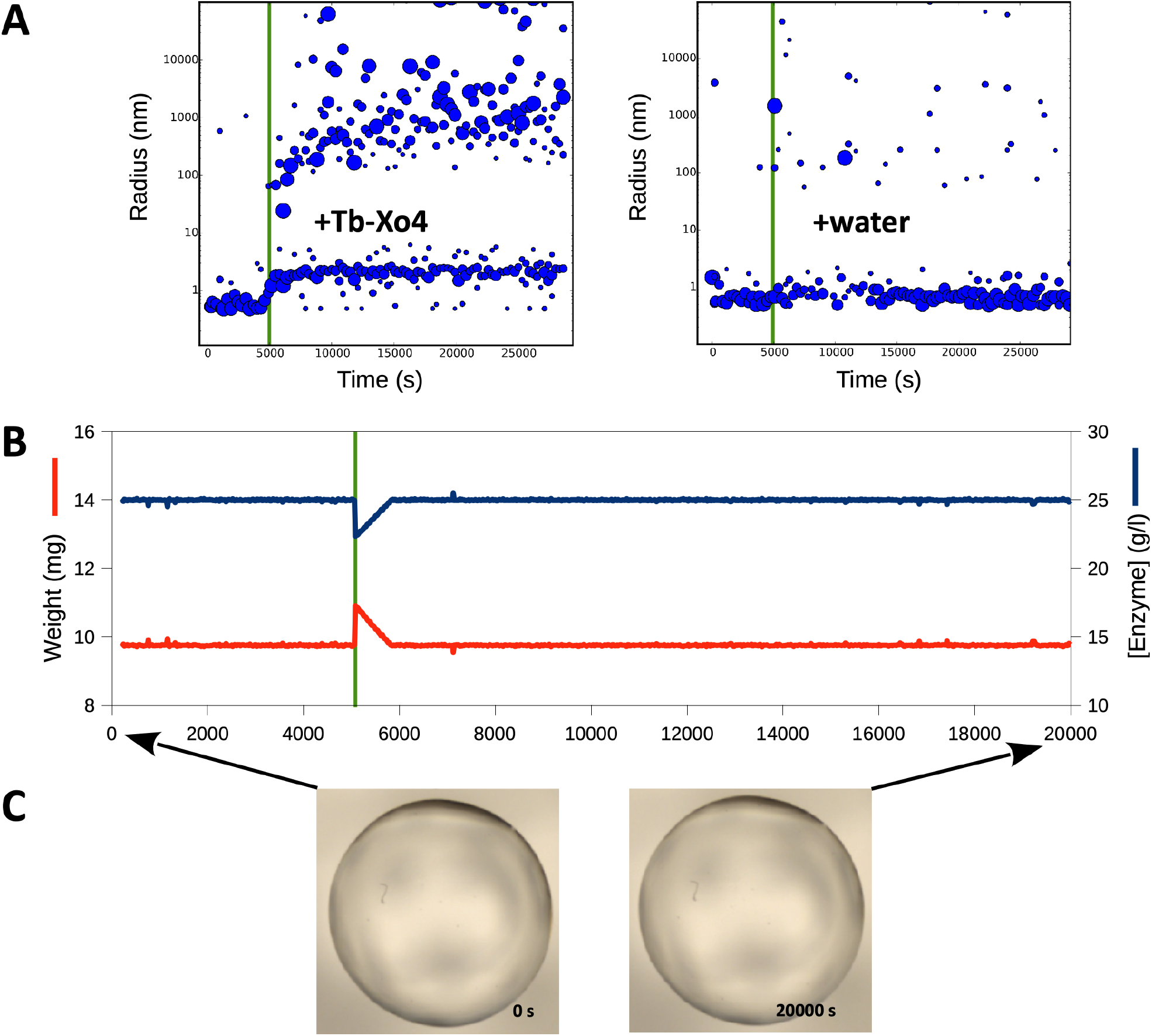
Monitoring the nucleant effect of Tb-Xo4 on HEWL. In this simple experiment, a Roche lysozyme solution (25 mg/ml in water) was incubated and maintained at a constant weight as illustrated by weight and concentration curves (B). Upon injection of 1 μl of nucleant (100 mM Tb-Xo4 solution), symbolized by the green line and highlighted by a 1 mg jump of the weight, the enzyme immediately reacted and formed larger objects that grew up to size of ~1 μm (A, left). After 8 h the drop remained clear (C) and was transferred to a Linbro plate for incubation. The control injection of 1 μl of water does not change the DLS signal (A, right).

**Figure 4** shows that the addition of Tb-Xo4 triggers the instantaneous formation of particles of larger size and this phenomenon of association, which is not observed upon water addition, leads to the growth of large crystals after weeks of incubation. Tb-Xo4 decreases the solubility of HEWL and promotes interactions between enzyme particles: the monomer seems to be converted to a slightly larger entity, possibly bridged by the nucleant (see **Figure 6B,C**) and large assemblies grow up to a size of 1 μm. Again, the XtalController was instrumental to highlight a situation leading to nucleation in a drop that remained clear for hours or days in videomicroscopy. Indeed, when a supersaturated state has been created, the system is thermodynamically set for either precipitation or nucleation. If nuclei are favored over precipitates then crystal growth is a matter of time (i.e. kinetics). The ‘activated’ drop is then simply transferred from the crystallization chamber to a Linbro plate for incubation until crystals grow

We further tested the nucleating property of Tb-Xo4 on the crystallization of HEWL from other suppliers (Sigma and Seikagaku) to check for potential enzyme preparation and batch effect. Following the protocol described in **Figure 4**, the concentration of enzyme was increased to 50 and 71 mg/ml to promote the growth of larger crystals. As for the Roche HEWL, the addition of Tb-Xo4 triggered the rapid formation (i.e. within 15 to 30 min) of populations of particles with radii ranging from 100 nm to 1000 nm in the XtalController (**Figure 5**, left panel). The same phenomenon was reproduced in the quartz cuvet of a Nanostar DLS instrument using the same lysozyme batch. Although they contribute significantly to the scattering signal, the percentage in mass of the larger populations was always low, in the range of a few percents (**Figure 5**, middle panel). In both cases, drops incubated for several months against a buffer reservoir produced crystals of several hundreds of μm suitable for crystallographic analyses.

**Figure 5:**
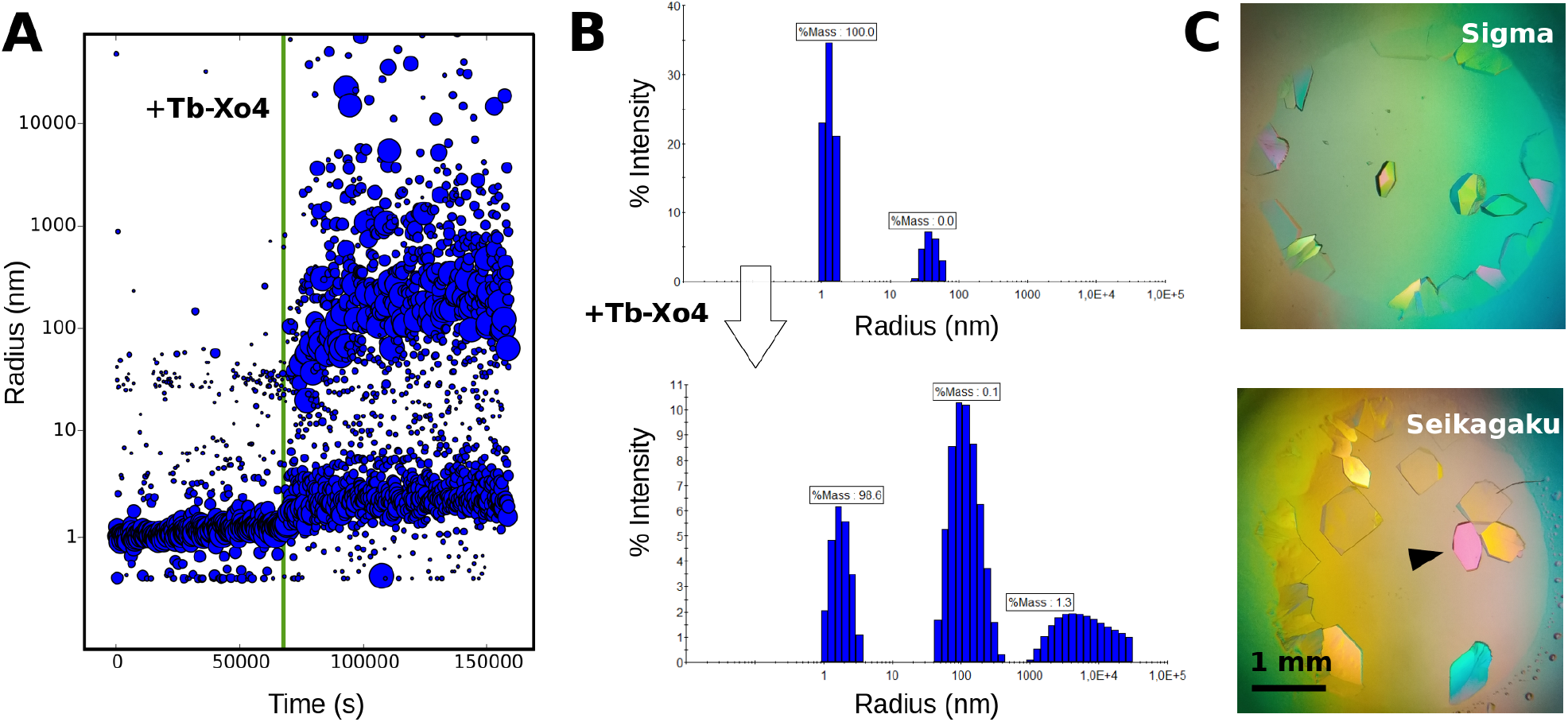
Production of Tb-Xo4-HEWL crystals for structural analyses. The experimental procedure described in **Figure 4** was applied to Sigma and Seikagaku lysozyme samples at a higher concentration (71 and 50 mg/ml, respectively) to promote the growth of larger crystals. A) the DLS monitoring of Sigma lysozyme in the XtalController upon Tb-Xo4 injection (green line) shows same features as in **Figure 4A**, yet amplified by the higher enzyme concentration. B) The phenomenon was followed on the same batch in a Nanostar DLS instrument. The top distribution detects almost exclusively the monomeric lysozyme before the injection, whereas populations with a *Rh* of ~100 nm and > 1 μm appear in the bottom distribution already 2.5 min after Tb-Xo4 injection. Crystallization assays performed with both batches were transferred from the XtalController to a Linbro plate for incubation. C) Images showing resulting crystals after 9 months before harvesting for data collection. The crystal marked by an arrow was used to collect the dataset presented in **Table 1**.

**Figure 6:**
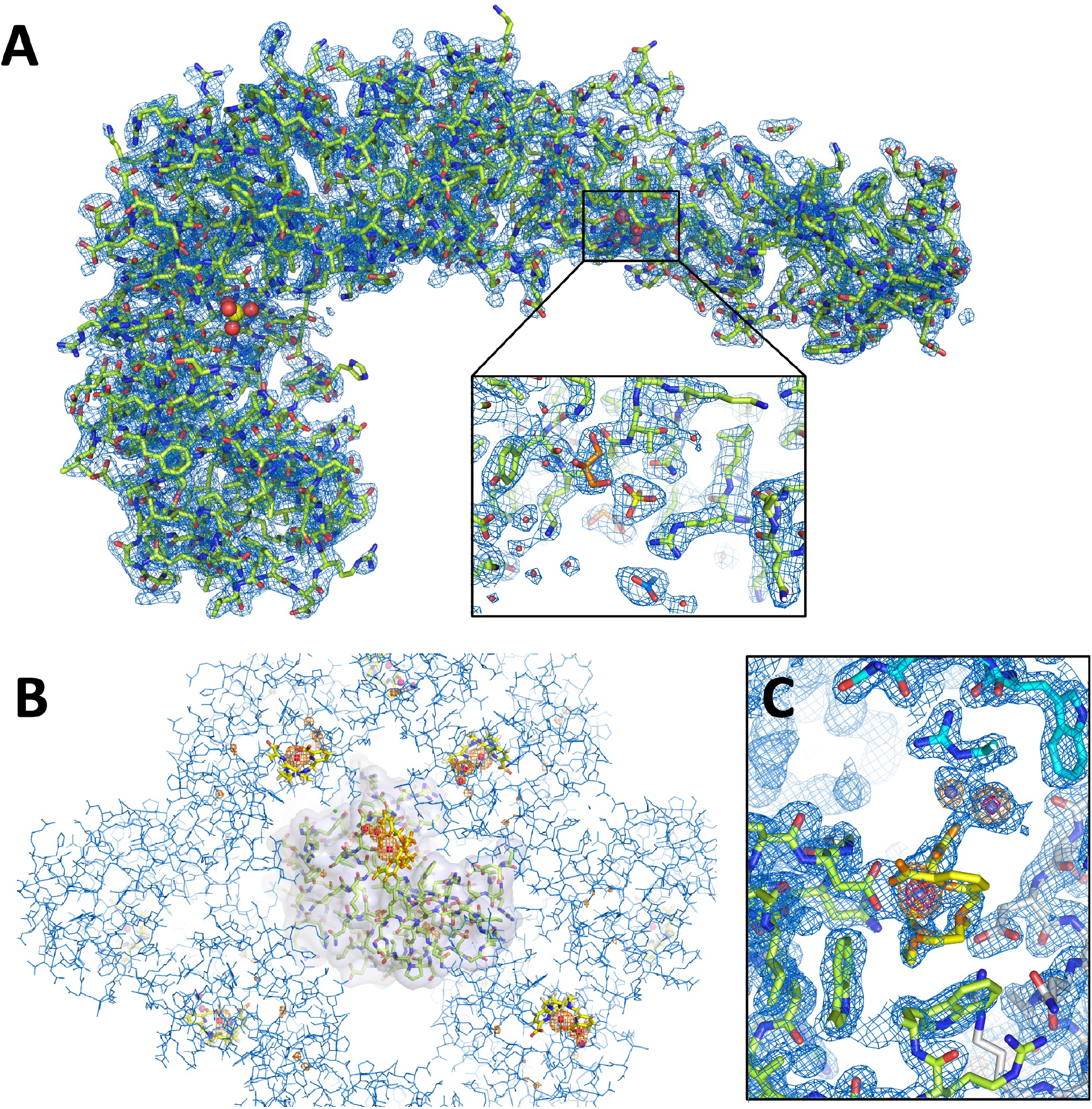
3D structures of PhaCCA and HEWL crystallized in the XtalController. A) Overall structure of PhaCCA and close-up on a phosphate, an acetate and two glycerol molecules bound to the enzyme surface. The blue *2mFo-DFc* map is contoured at 1.2 σ. B) View of tetragonal crystal packing of HEWL seen down the 4-fold axis. One lysozyme monomer is represented with a Tb-Xo4 complex bound to its surface. Symmetry related monomers are depicted in line mode. The anomalous difference map shown in orange is contoured at 5 σ and indicates the presence of Tb^3+^ ions (red spheres) as well as sulfur atoms. C) Zoom on the Tb-Xo4 binding sites. The side chain of Asp101 directly coordinates the Tb^3+^ of the major site for which the organic ligand (in yellow) is partially visible in the blue *2mFo-DFc* map contoured at 1 σ. The anomalous difference map depicted in orange and red is contoured at 5 and 20 σ, respectively, and highlights the strong anomalous signal of Tb^3+^ ions. Two alternate positions are observed but their ligand is not visible due to low occupancy. The two adjacent Tb-Xo4 sites constitute a bridge (molecular glue) between the green and blue lysozyme monomers.

### 3.3 Characterization of crystals grown in the XtalController

The quality of PhaCCA and HEWL crystals grown using the XtalController was assessed by X-ray diffraction. Crystals of PhaCCA were analyzed under cryogenic conditions at the FIP-BM30A synchrotron beamline (ESRF, France). **Table 1** shows excellent statistics for such a crystal leading to a complete diffraction dataset at 2.28 Å resolution. The signal extended isotropically up to 1.8 Å (not shown) as for our best data collected on a much stronger beamline installed on an insertion device (PROXIMA2A, SOLEIL, France). Here, data were truncated because of the presence of two diffraction rings at resolutions of about 1.9 and 2.15 Å. They were probably due to suboptimal cryocooling of the solvent and significantly altered the statistics in the high resolution shell (<2.2 Å). However, the resulting electron density maps are extremely clear and allowed to observed two phosphate ions (**Figure 6A**) from the crystallant bound to the surface of this RNA binding enzyme.

**Table 1.** Statistics of crystal analysis and structure refinement.

| **Enzyme** | *PhaCCA* | *HEWL* |
| --- | --- | --- |
| **Data collection** |  |  |
| X-ray source | FIP/BM30A – ESRF | Rigaku Fr-X – IGBMC |
| Wavelength (Å) | 0.9799 | 1.5418 |
| Detector | ADSC Q315r | EIGER R 4M |
| Temperature (K) | 100 | 293 |
| Space group | $P4_32_12$ | $P4_32_12$ |
| Cell parameters (Å) | 70.53, 70.53, 291.48 | 78.81, 78.81, 38.33 |
| Mosaicity (°) | 0.31 | 0.12 |
| Solvent content | 67.4 | 40.8 |
| Resolution range (Å) | 2.28 – 47 (2.28 – 2.42) | 1.51 – 35 (1.51 – 1.60) |
| Number of reflections | 237664 (37915) | 164648 (14601) |
| Number of unique reflections | 34470 (5385) | 19535 (3017) |
| Multiplicity | 6.9 (7.0) | 8.4 (4.8) |
| Completeness (%) | 99.5 (98.5) | 99.2 (96.5) |
| Mean I/sigma(I) | 15.5 (1.5) | 29.7 (1.9) |
| R<sub>meas</sub> (%) | 8.6 (126) | 3.6 (72.6) |
| SigAno | - | 2.1 (0.6) |
| CC<sub>1/2</sub> | 99.9 (76.0) | 100 (78.1) |
| **Structure refinement** |  |  |
| R<sub>work</sub>, R<sub>free</sub> | 0.212, 0.255 | 0.143, 0.177 |
| Number of non-H atoms |  |  |
| enzyme, ligands, solvent | 2989, 40, 105 | 1001, 35, 54 |
| RMSD on bonds (Å) and angles (°) | 0.009, 0.96 | 0.004, 0.69 |
| Average ADPs (Å<sup>2</sup>) |  |  |
| overall, enzyme, ligands, solvent | 52.9, 52.7, 65.7, 52.8 | 30.6, 29.1, 56.6, 42.2 |
| Ramachandran plot: % of residues in |  |  |
| favored, allowed, unfavored regions | 96.4, 3.3, 0.3 | 99.2, 0.8, 0 |

Lysozyme crystals were analyzed at room temperature using the X-ray lab source from the INSTRUCT-FRISBI platform at the IGBMC (Illkirch, France). A fast data collection was applied using an EIGER pixel detector to collect complete anomalous data before crystal decay. The Tb2+ ions give a strong anomalous signal (f’’ of 9 electrons) at the Cu Kα wavelength (1.5418 Å). The dataset collected at 1.51 Å resolution (**Table 1**) allowed the identification of three Tb^3+^ sites, a major one for which the ligand could be built, and two minor sites (**Figure 6C**). These sites are consistent with those described in PDBid 6F2I determined with a HEWL crystal grown in 0.8 M NaCl and 100 mM Tb-Xo4 [20]. The low occupancy of Tb^3+^ sites and the absence of visible ligand for the two minor sites are also consistent with the low concentration of Tb-Xo4 (10 mM) which provide a ratio of two Tb-Xo4 complexes for one lysozyme molecule. This concentration seems to be sufficient to trigger the association of lysozyme particles, nucleation and subsequent crystal growth, but not to guarantee a full occupancy in the crystal lattice.

## 4. Conclusion

In the perspective of emerging time-resolved studies of enzyme:substrate systems using SFX and XFEL facilities, it becomes increasingly important to gain more control over sample production, quality and reproducibility. These two examples of enzyme crystallization highlight the usefulness of the XtalController to monitor the evolution of crystallization assays and to act on the process. Beyond helping define and optimize crystallization conditions, the XtalController and its integrated DLS module may also be an ideal tool to:

- explore phase diagrams of biomolecules with a direct feedback on nucleation events,
- study the stability of biomolecules in solution with respect to various parameters such as temperature, pH, ligands, etc.,
- determine the optimum conditions for introducing a cryoprotectant,
- ensure the reproducibility of crystals in the context of structural biology investigations,
- produce calibrated nanocrystals on demand (difficult to monitor and control otherwise) for diffraction analyses using X-ray free electron lasers and CryoEM, or, conversely, to promote the selective growth of large crystals for neutron diffraction.

More generally, this type of versatile instruments provides a more rational approach to crystallization and a great alternative to extensive blind screening. We do believe that this technology has a bright future.

## Acknowledgements

The authors thank the team of FIP beamline at the European Synchrotron Radiation Facility (ESRF, Grenoble, France) for beamtime allocation, Alexandra Bluhm and Léna Coudray for their contribution to crystallization experiments and structure refinement, Robin Schubert and Christian Betzel (University of Hamburg), as well as Karsten Dierks, Arne Mayer and Annette Eckhardt (Xtal-Concepts GmbH, Hamburg) for fruitful discussions and exchanges about the XtalController.

## Author Contributions

R.dW., K.R., O.H., B.L., A.McE., F.R., O.M., E.G., S.E. and C.S. designed and performed the experiments. R.dW., K.R., B.L., P.B. and C.S. analyzed the data. All authors helped with writing the manuscript.

